# A novel role for TRPV1 in macrophage giant cell formation

**DOI:** 10.64898/2026.05.11.724406

**Authors:** Karunakaran R. Sankaran, Mohammad I. Khan, Shaik O. Rahaman

## Abstract

TRPV1 (transient receptor potential vanilloid 1) is a non-selective cation channel with high permeability to Ca^2+^ and is best known for its roles in sensory signaling. However, its function in immune cell biology, particularly in macrophage fusion, remains unknown. Cell fusion is a critical process in both physiological and pathological contexts, including development, tissue remodeling, and the foreign body response (FBR) to implanted biomaterials. During FBR, macrophages undergo fusion to form multinucleated foreign body giant cells (FBGCs), which contribute to implant degradation and fibrotic encapsulation. Here, we identify TRPV1 as a key regulator of macrophage multinucleation and FBGC formation. We demonstrate that TRPV1 is endogenously expressed in bone marrow-derived macrophages (BMDMs) and is upregulated in response to fusogenic cytokines and inflammatory stimuli. Functionally, TRPV1 promotes matrix stiffness-dependent macrophage adhesion and spreading, indicating a role in mechanosensitive signaling. We show that TRPV1 is required for efficient macrophage fusion under both cytokine-driven and matrix stiffness-mediated conditions. Mechanistically, TRPV1 links extracellular mechanical cues and cytokine signaling to cytoskeletal remodeling, facilitating the actin reorganization necessary for cell fusion. Importantly, TRPV1 deficiency does not alter TRPV4-mediated Ca^2+^ signaling, demonstrating that TRPV1 operates independently of TRPV4, a known mechanosensitive channel implicated in FBR and FBGC formation. Collectively, these findings suggest TRPV1 as a previously unrecognized mechanosensitive regulator of macrophage fusion and FBGC formation. This work provides new insight into the molecular mechanisms governing FBR and identifies TRPV1 as a potential therapeutic target for improving biomaterial biocompatibility and mitigating fibrosis.

## Introduction

TRPV1, originally identified as the capsaicin receptor, is a non-selective cation channel that conducts several ions, including H^+^, Na^+^, Ca^2+^, and Mg^2+^, with particularly high permeability to Ca^2+^ (1-3). The TRPV1 protein consists of 838 amino acids. Structurally, TRPV1 forms a tetrameric channel in which each subunit contains six transmembrane domains, with a pore region located between the fifth and sixth segments. Both the N-terminal and C-terminal domains are positioned on the cytoplasmic side of the membrane and play important roles in regulating channel activity (4). Functionally, TRPV1 acts as a polymodal nociceptive receptor that can be activated by a wide range of soluble and physical stimuli, including capsaicin, lipopolysaccharide, hyperthermia, acidic pH, reduced osmolarity, and various endogenous or synthetic ligands (1-6). While TRPV1 is predominantly associated with sensory neurons in the nervous system, it is also expressed in multiple peripheral tissues such as the heart, lungs, kidneys, skin, intestines, and bladder (7-14). At the cellular level, TRPV1 has been detected in both neuronal and non-neuronal cell types, including lymphocytes, macrophages, dendritic cells, mast cells, endothelial cells, vascular smooth muscle cells, epithelial cells, and fibroblasts (15-22).

TRPV1 channels permit Ca^2+^ influx and therefore participate in calcium-dependent signaling pathways that regulate numerous cellular functions, including proliferation, survival, migration, embryonic development, apoptosis, inflammatory responses, nociception, and tumor metastasis (23-28). Activation of TRPV1 increases membrane permeability to Na^+^ and Ca^2+^, promoting ion entry and triggering downstream signaling cascades. Among these pathways, TRPV1 activation can induce transactivation of the epidermal growth factor receptor (EGFR), which subsequently stimulates MAPK and Akt/PI3K signaling in human corneal epithelial cells (29). In addition, TRPV1 signaling contributes to transforming growth factor-β1 (TGF-β1)-driven differentiation of stromal keratocytes and renal mesenchymal cells into myofibroblasts, processes associated with corneal opacity and renal fibrosis (30, 31).

Activation mechanisms of nociceptive TRPV channels have been extensively characterized in sensory neurons, where they function as key mediators of pain perception and neurogenic inflammation (20, 32, 33). In neuronal systems, these channels are activated by diverse physical and soluble/chemical stimuli and initiate Ca^2+^-dependent signaling cascades that regulate neuronal excitability and neurotransmitter release. In contrast, the functional significance and activation pathways of TRPV channels in immune cells remain incompletely understood. Although TRPV family members, including TRPV1, have been detected in several immune cell types, their precise physiological roles in immune regulation are still being elucidated. Moreover, findings from different studies have often been inconsistent or contradictory, with some reports suggesting pro-inflammatory functions while others indicate anti-inflammatory or immunomodulatory roles. These discrepancies likely reflect differences in experimental models, cell types, microenvironmental conditions, pathophysiological condition, and signaling contexts. Consequently, further investigation is needed to clarify how TRPV1 activation influences immune cell behavior and how these pathways contribute to inflammation, cell differentiation, tissue remodeling, and disease progression.

Evidence from genetic and pharmacological studies further highlights the role of TRPV1 in tissue remodeling and fibrotic disease. Deletion of TRPV1 suppresses osteoclast formation and differentiation in TRPV1-deficient mice (34, 35). Consistently, the TRPV1 antagonist capsazepine inhibits osteoclast differentiation in vitro and reduces ovariectomy-induced bone loss in vivo (36). Studies employing TRPV1 agonists, antagonists, and knockout models collectively demonstrate a strong link between TRPV1 activity and the progression of fibrotic disorders (37-42). For example, TRPV1 has been implicated in the regulation of fibrotic processes affecting the kidney, lungs, heart, and cornea (42-44). Given the shared mechanisms underlying many fibrotic diseases, clarifying both the direct and indirect contributions of TRPV1 to fibrosis may provide important insights into disease pathogenesis and support the development of new therapeutic strategies.

In this work, we show that TRPV1 is a mechanosensitive regulator of macrophage fusion, driving matrix stiffness- and cytokine-induced multinucleation through cytoskeletal remodeling. It functions independently of TRPV4, identifying TRPV1 as a novel target to modulate FBGC formation, FBR, and biomaterial-associated fibrosis.

## Materials and methods

### Materials

Dulbecco’s modified Eagle’s medium (DMEM), heat-inactivated fetal bovine serum (FBS), and routine cell culture supplies were obtained from Gibco. Antibiotic-antimycotic (A/A) solution, lipopolysaccharide (LPS; cat# L5418), the TRPV1 agonist capsaicin, the TRPV1 antagonist AMG, and the TRPV2 inhibitor SKF96365 were purchased from Sigma-Aldrich (St. Louis, MO). ProLong™ Diamond mounting medium with 4′,6-diamidino-2-phenylindole (DAPI) was obtained from Thermo Fisher Scientific. Recombinant cytokines, including interleukin-4 (IL-4), granulocyte-macrophage colony-stimulating factor (GMCSF), and macrophage colony-stimulating factor (MCSF), were sourced from R&D Systems. Primary antibodies comprised anti-actin (cat# 4970S; Cell Signaling Technology, Danvers, MA) and anti-TRPV1 (cat# ACC-030), anti-TRPV2 (cat# ACC-039), and anti-TRPA1 (cat# ACC-037) from Alomone Labs. Species-appropriate secondary antibodies (goat, rabbit, and mouse) were obtained from Jackson ImmunoResearch. The TRPV4 agonist GSK1016790A (GSK101; cat# 6433) was purchased from Tocris. Petrisoft plates and collagen-coated polyacrylamide (PA) hydrogels with defined stiffness (1 kPa, cat# PS100-COL-1-EA; 50 kPa, cat# PS100-COL-50-EA) were obtained from Matrigen.

### Animal maintenance and cell culture

C57BL/6 and TRPV1 KO mice were purchased from The Jackson Laboratory (ME, USA). All animal experiments were conducted with protocols approved by the Institutional Animal Care and Use Committee (IACUC) at the University of Maryland College Park (Protocol No. R-OCT-24-36). Animals were housed in a pathogen-free environment under controlled temperature and humidity conditions, with food and water provided ad libitum. Bone marrow was harvested from mouse femur and incubated with 25 ng/ml MCSF supplemented DMEM (10% FBS; 1x A/A) for 7-8 days to obtain mature bone marrow-derived macrophages (BMDMs) (21, 24, 26).

### Immunoblotting

For Western blot analysis to detect the levels of TRPV1, TRPV2, TRPA1, and actin levels, WT BMDMs were either treated with LPS (50 and 250 ng/ml) or IL4 plus GMCSF (25 and 50 ng/ml) for 48 h or kept untreated for the control group. Whole cell extracts were separated using 10 % SDS-PAGE and probed with antibodies against TRPV1, TRPV2, TRPA1, and actin. The blots were visualized using HRP-conjugated secondary IgGs and analyzed by UVP BioSpectrum.

### Calcium influx measurement by spinning disc confocal microscopy

Fluorescence imaging of macrophages was done using PerkinElmer UltraView VoX confocal spinning disk microscope equipped with Nikon TI inverted stand and 20 x objective lens. WT and TRPV1 KO macrophages (∼ 5 × 10^5^) were cultured on 35 mm glass bottom plates in DMEM supplemented with 10% FBS and 1x A/A. After 48h of incubation, medium was replaced with 1% BSA containing DMEM and 1x A/A. Then cells were treated with 25 ng/mL of IL4/GMCSF and incubated for 48 h at 37°C. For the Ca^2+^ influx measurement, the cells were incubated with 5 μM Calbryte 590 AM dye in 1× HBSS buffer with 1mM probenecid (Millipore Sigma) for 1 h at 37°C. Transient fluorescence measurement was performed by perfusing the cells with or without capsaicin (10 μM) in HBSS buffer for 2 mins in the dark and fluorescence was measured for 10 mins. The fluorescence intensity was analyzed and quantified using Image J. The imaging was conducted at 1.57 frames per second. The ROI was determined separately for each cell. For each frame, cells in the control untreated condition were outlined using the freeform tool in ImageJ. Intensity values were extracted using the ROI manager. The same ROIs were applied to the corresponding frames in treated conditions. Intracellular Ca^2+^ dynamics were assessed using the FLIPR Calcium 6 assay kit (Molecular device; Explorer Kit, PN: R8190). Around 2 × 10^5^ TRPV1 KO BMDMs were plated in 96-well plates (Greiner Sensoplate glass bottom) in 200 μL DMEM (10% FBS and 1x A/A) and cultured for 24 h. Then media was replaced with 200 μL of DMEM (1% BSA and 1x A/A) in the presence or absence of 25 ng/mL IL-4 and GMCSF, and incubated for 48 h at 37°C. Then 100 µL media was discarded and 100 µL of Calcium 6 dye prepared in 1× HBSS supplemented with 20 mM HEPES (pH 7.4), and 2.5 mM probenecid was added into each well followed by incubation for 60 min at 37°C. Plates were transferred to a FlexStation 3 system for real-time fluorescence measurements. TRPV4 activation was induced by addition of its selective agonist GSK 1016790A (100 nM) and Ca^2+^ influx was measured for 180 seconds and quantified as changes in fluorescence intensity (ΔF/F, Max-Min). Results are presented as relative fluorescence units (RFU) (45-47).

### Adhesion and spreading of BMDMs on polyacrylamide hydrogels

Mature BMDMs were placed on coverslips and collagen-coated (10 μg/ml) PA hydrogels with stiffness levels of 1 kPa and 50 kPa. This culture was maintained in DMEM supplemented with 10% FBS. Subsequently, the cells were subjected to treatment, either with or without IL4 plus GMCSF (25 ng/ml), for a duration of 24 hours before being examined under a microscope.

### Macrophage giant cell formation

Bone marrow cells were harvested from femurs of wild-type (WT) and TRPV1 KO mice and differentiated into BMDMs in DMEM containing 10% FBS and 25 ng/mL MCSF at 37°C in a 5% CO_2_ atmosphere for 7-8 days (48-50). Fully differentiated BMDMs were subsequently plated on permanox culture dishes or collagen-coated PA hydrogels of defined stiffness (1 or 50 kPa). To induce foreign body giant cell (FBGC) formation, cells were treated with IL-4 and GMCSF (25 ng/mL each) every 48 h for 4-6 days until maximal fusion was achieved (51-53). Cultures were stained with Giemsa, and FBGC number, average size per field, and total FBGC area were quantified. Fusion efficiency was determined as the proportion of nuclei within multinucleated giant cells (> 5 nuclei per cell) relative to the total nuclei count (51-53).

### Transfection of BMDMs

For FBGC formation studies, 4 × 10^5^ BMDMs were plated onto Permanox 8-well chamber slides in DMEM supplemented with 10% FBS and allowed to adhere for 24 h at 37°C in 5% CO_2_. Cells were then transfected with control siRNA (TRPV1-Scr, 50 nM) or TRPV1-targeting siRNA (10 or 50 nM). Following a 48 h incubation under standard culture conditions, macrophages were stimulated with IL-4 and GM-CSF (25 ng/mL each) for an additional 48 h to promote multinucleated giant cell formation. Cells were subsequently fixed in 10% formalin, stained with Giemsa, and imaged using a Nikon Eclipse microscope.

### Fluorescence microscopy

For immunofluorescence analysis, BMDMs were cultured on Permanox slides and stimulated with IL-4 and GMCSF (25 ng/mL each) for 7-8 days. Cells were fixed in 3% paraformaldehyde, permeabilized using 0.1% Triton X-100, and blocked with 1% BSA for 1 h. Samples were then incubated overnight at 4°C with primary antibodies against TRPV1. Following washing, appropriate Alexa Fluor-conjugated secondary antibodies were applied. F-actin was visualized using Alexa Fluor-conjugated phalloidin, and nuclei were counterstained with DAPI. Images were acquired using a Zeiss Axio Observer Z1 fluorescence microscope.

### Statistical analysis

All data are presented as mean ± SEM. Statistical comparisons between different groups were conducted using Student’s t-test or one-way ANOVA followed by the Bonferroni test. In the notation, “ns” indicates not significant, * denotes p < 0.05, ** denotes p < 0.01, *** denotes p < 0.001, and **** denotes p < 0.0001.

### Declaration of generative AI and AI-assisted technologies in the writing process

During the preparation of this work the author(s) used ChatGPT 5.2 in order to improve readability and perform proofreading. After using this tool/service, the author(s) reviewed and edited the content as needed and take(s) full responsibility for the content of the publication.

## Results

### Fusogenic cytokines and LPS treatment induce TRPV1 expression in BMDMs

Cell fusion is a fundamental process underlying diverse physiological and pathological events, including fertilization, development, bone remodeling, and host responses to pathogens and implanted materials (54-56). Macrophages are key mediators of the FBR to biomaterials and medical devices, contributing to the formation of multinucleated FBGCs that participate in implant degradation, as well as to fibrotic encapsulation of implants (48-53, 57-60). Despite growing interest in TRPV channels in immune regulation, their functional roles and activation mechanisms in macrophages remain poorly defined. Although several TRPV family members, including TRPV1, have been identified in immune cells, their involvement in macrophage fusion and FBGC formation is not determined. To assess TRPV1 expression, we performed immunoblotting and immunofluorescence analyses in BMDMs. These studies revealed endogenous expression of both TRPV1 and TRPA1 (Figure 1A-C, 1D-E). Notably, stimulation of BMDMs with fusogenic IL-4 plus GMCSF, cytokines known to promote FBGC formation, or with LPS, a pro-inflammatory stimulus, led to 4-fold increased TRPV1 expression (Figure 1A-B, 1D-E). Collectively, these findings demonstrate that TRPV1 is expressed in BMDMs and is upregulated in response to both fusogenic cytokine signaling and inflammatory stimulation, implicating a potential role for TRPV1 in macrophage activation and fusion.

**Figure 1.**
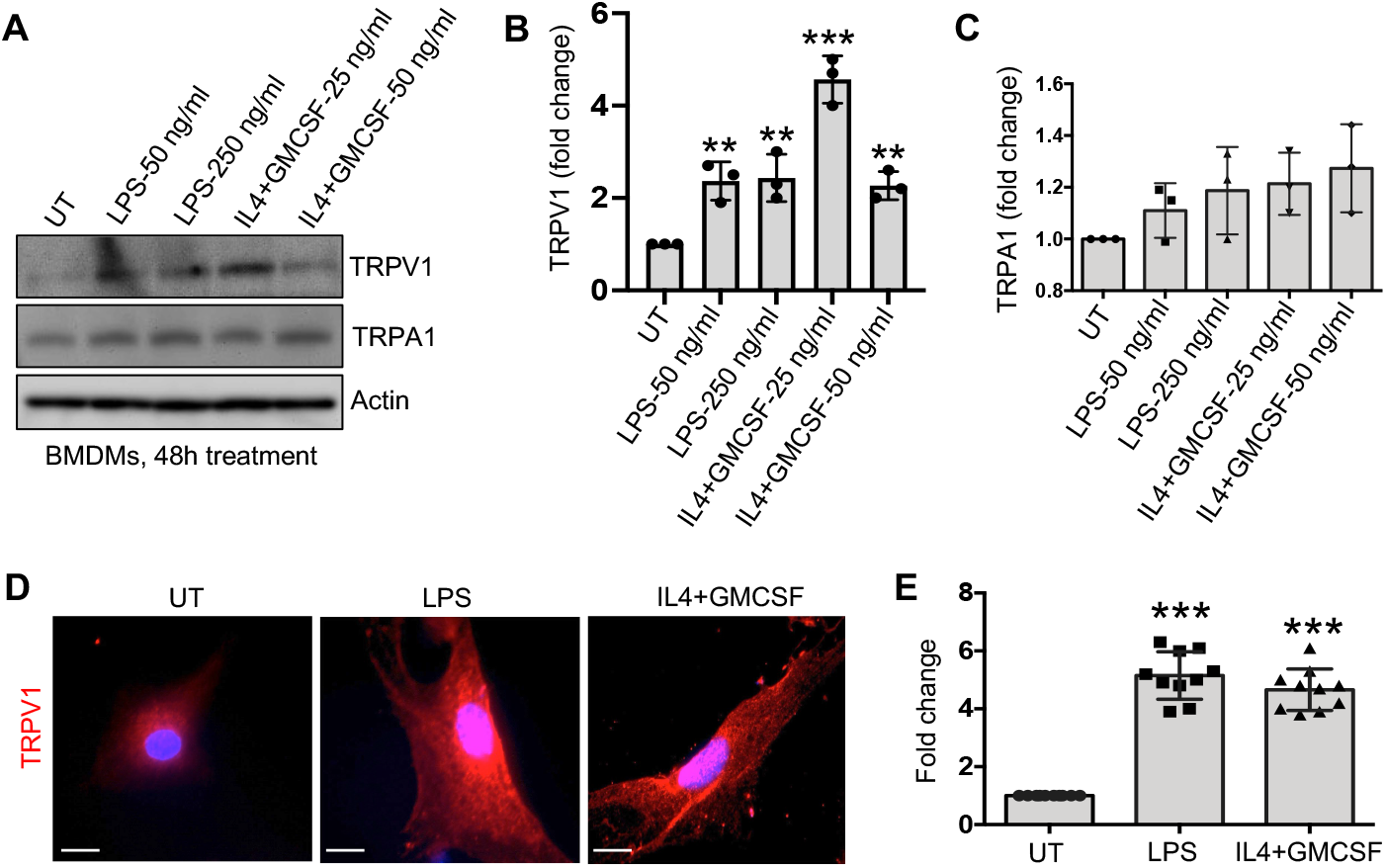
Fusogenic cytokines and LPS upregulate TRPV1 expression in BMDMs. (A) Immunoblot analysis shows TRPV1, TRPA1, and actin levels in WT BMDMs 48 h following treatment with or without IL-4 plus GMCSF (25 ng/ml). (B-C) Densitometric quantification of immunoblot data from (A) (n = 3 biological replicates; one-way ANOVA, **p < 0.01, ***p < 0.001). (D) Representative immunofluorescence images of WT BMDMs stained for TRPV1 (red) using anti-TRPV1 IgG (original magnification, 60 x; scale bar, 2 μm; n = 10 cells per condition). Statistical analysis by Student’s t-test, ***p < 0.001.

### TRPV1 mediates matrix stiffness-induced adhesion and spreading in BMDMs

Emerging studies highlight extracellular matrix stiffness as a key regulator of fundamental cellular behaviors (49, 51-53, 61-70). Consistent with prior work from our group and others, macrophage activities including differentiation, adhesion, migration, and proliferation are strongly influenced by the mechanical properties of their environment (49-53). Cells adapt to external rigidity by remodeling cytoskeletal tension, enabling effective force generation during migration (71-75). To examine whether TRPV1 contributes to macrophage adhesion and spreading-processes central to migration, we cultured WT BMDMs on collagen-coated (10 µg/ml) polyacrylamide hydrogels spanning physiologically relevant stiffnesses: soft (1 kPa) and stiff (50 kPa). These conditions approximate normal tissue compliance (∼1 kPa) and fibrotic states (∼25-50 kPa). Cells were treated with or without the TRPV1 antagonist AMG9810 [(2E)-N-(2,3-dihydro-1,4-benzodioxin-6-yl)-3-[4-(1,1-dimethylethyl)phenyl]-2-propenamide], in the presence or absence of IL-4 plus GMCSF stimulation. We observed enhanced adhesion and spreading of BMDMs on stiff substrates relative to soft matrices (Figure 2A-D). Cytokine priming with IL-4 and GM-CSF further amplified these responses. Notably, inhibition of TRPV1 with AMG9810 abolished cytokine-driven increases in adhesion and spreading, supporting a central role for TRPV1 in mechanosensitive regulation of macrophage behavior under physiologically relevant conditions.

**Figure 2.**
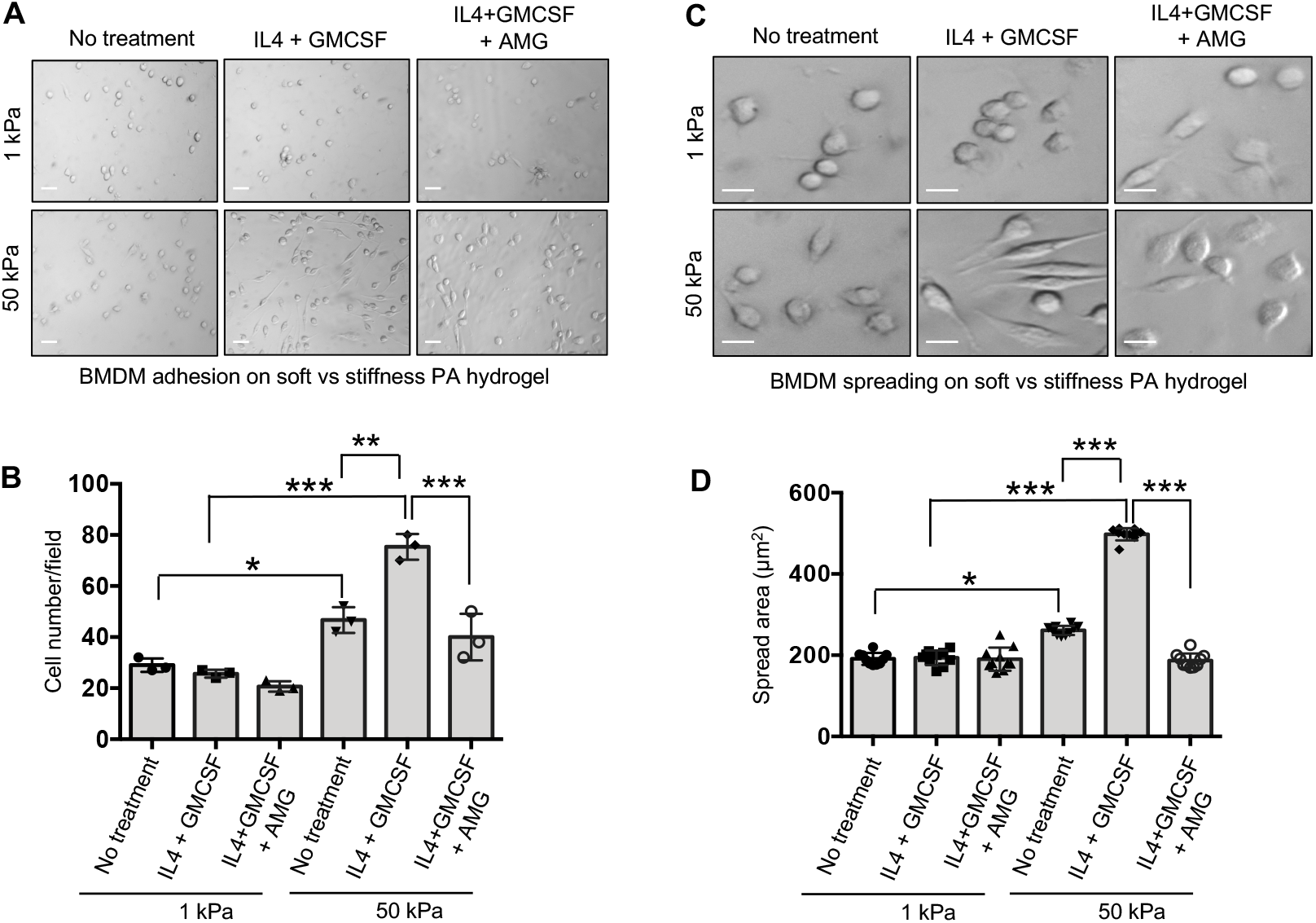
TRPV1 regulates stiffness-dependent adhesion and spreading in BMDMs. (A) Representative images of BMDMs cultured on collagen-coated (10 μg/ml) PA hydrogels of defined stiffness (1 kPa and 50 kPa), showing cell adhesion under AMG-treated or untreated conditions, with or without IL-4 plus GMCSF stimulation (25 ng/ml, 24 h). Scale bar, 50 μm. (B) Quantification of adherent cells per field (n = 3 fields per condition). (C) Representative images illustrating cell spreading of BMDMs plated on 1 kPa and 50 kPa hydrogels for 48 h under AMG-treated or untreated conditions, in the presence or absence of IL-4 plus GMCSF (25 ng/ml). Scale bar, 10 μm. (D) Quantification of mean cell spread area (n = 10 cells per condition). Statistical analysis by one-way ANOVA, *p < 0.05, **p < 0.01, ***p < 0.001.

### TRPV1 participates in multinucleated macrophage giant cell formation

Macrophages are central to the FBR, driving both multinucleated FBGC formation and fibrotic encapsulation of implanted materials (48-53, 57-60). While TRPV channels have been increasingly implicated in immune cell function, their specific contribution to macrophage fusion remains unclear. To address this, we employed an established IL-4 plus GMCSF-driven *in vitro* FBGC formation model to evaluate the role of TRPV1. Pharmacological inhibition of TRPV1 with AMG9810 markedly attenuated macrophage fusion compared to untreated controls or cells exposed to the TRPV2 inhibitor SKF96365. Quantitative analysis revealed substantial reductions in FBGC number (∼6-fold), fusion index (∼12-fold), and FBGC size (∼4-fold) (Figure 3A-D), indicating a strong dependence on TRPV1 activity. To independently validate these findings, TRPV1 expression was silenced in BMDMs using targeted siRNA. Both immunofluorescence and immunoblotting confirmed efficient knockdown (> 80%) relative to scramble controls (Figure 3E-G). Consistent with the pharmacological data, TRPV1-deficient macrophages exhibited pronounced defects in fusion, with decreases in FBGC formation (∼5-fold), fusion index (∼5-fold), and cell size (∼4-fold) (Figure 3H-J). Together, these results establish TRPV1 as a critical regulator of macrophage multinucleation and FBGC formation.

**Figure 3.**
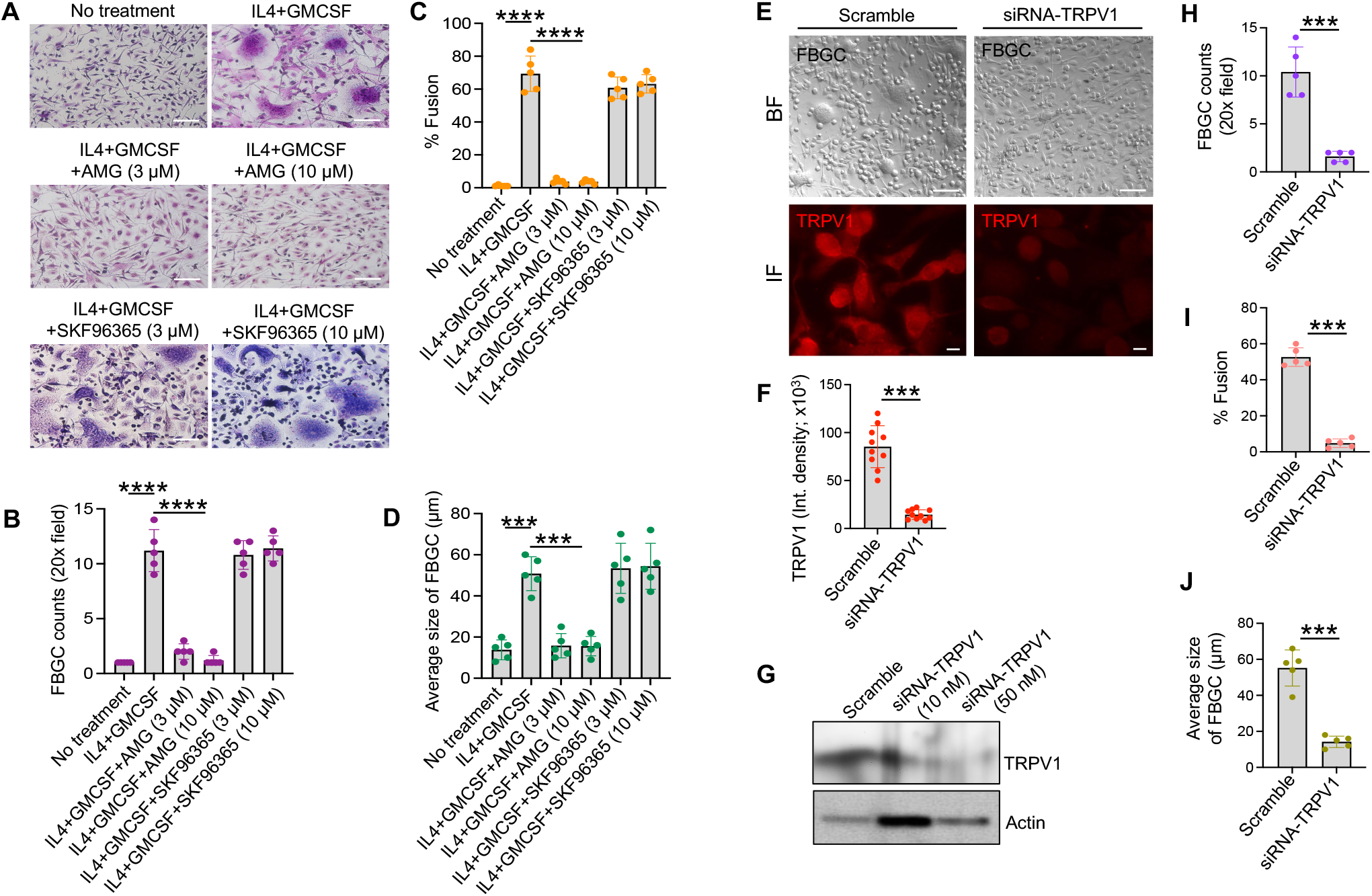
TRPV1 contributes to multinucleated FBGC formation in macrophages. (A) Representative Giemsa-stained images of multinucleated FBGCs in WT BMDMs left untreated or stimulated with IL-4 plus GMCSF (25 ng/ml, 96 h), in the presence or absence of the TRPV1 antagonist AMG. (B-D) Quantitative analysis of FBGC formation from (A): (B) number of FBGCs per high-power field, (C) percentage of fused BMDMs, and (D) average FBGC size. Data represent n = 3 biological replicates with 5 images per group; scale bar, 100 μm; Student’s t-test, ***p < 0.001, ****p < 0.0001. (E) Representative immunofluorescence images of WT BMDMs transfected with scramble or TRPV1-targeting siRNA, stained for TRPV1 (red) using anti-TRPV1 IgG (original magnification, 60x; scale bar, 2 μm). (F) Quantification of TRPV1 fluorescence intensity (n = 10 cells per condition; Student’s t-test, ***p < 0.001). (G) Immunoblot showing TRPV1 expression in WT BMDMs 48 h after transfection with scramble or TRPV1 siRNA. (H-J) Quantification of FBGC formation following TRPV1 knockdown: (H) number of FBGCs per high-power field, (I) percentage of fused BMDMs, and (J) average FBGC size. Data represent n = 3 biological replicates with 5 images per group; Student’s t-test, ***p < 0.001.

### TRPV1 regulates matrix stiffness-induced macrophage giant cell formation

Matrix stiffness is increasingly recognized as key determinants of the FBR, shaping macrophage behavior and FBGC formation (49-53). To investigate whether TRPV1 mediates matrix stiffness-dependent macrophage fusion, WT BMDMs were cultured on collagen-coated PA hydrogels spanning physiologically relevant compliance (1 kPa) to fibrotic-like stiffness (50 kPa). FBGC formation was enhanced on stiffer substrates compared to compliant matrices, and this effect was further potentiated by IL-4 and GMCSF stimulation (Figure 4A-C), indicating that both mechanical and cytokine cues cooperatively drive macrophage fusion. Pharmacological inhibition of TRPV1 with AMG9810 abrogated cytokine-induced increases in FBGC formation across stiffness conditions, implicating TRPV1 in the integration of biochemical and mechanical signals.

**Figure 4.**
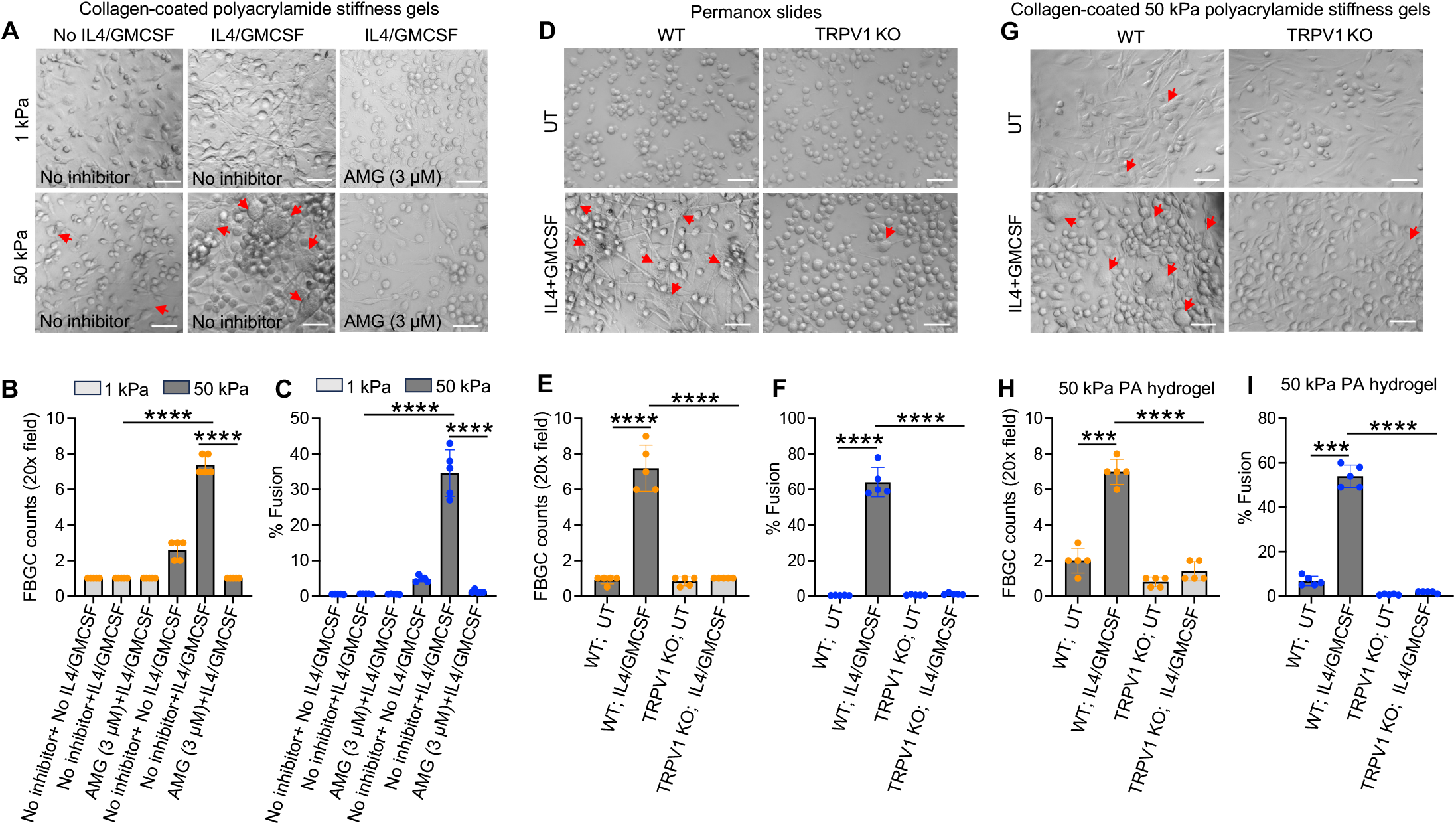
TRPV1 regulates matrix stiffness-dependent macrophage giant cell formation. (A) Representative images of FBGCs formed by WT BMDMs cultured on collagen-coated (10 μg/ml) PA hydrogels of defined stiffness (1 kPa and 50 kPa), left untreated or stimulated with IL-4 plus GMCSF (25 ng/ml, 96 h), in the presence or absence of the TRPV1 antagonist AMG. (B-C) Quantification of FBGC formation from (A): (B) number of FBGCs per high-power field and (C) percentage of fused BMDMs. (D) Representative images of FBGCs in WT and TRPV1 KO BMDMs cultured on Permanox slides under untreated or IL-4 plus GMCSF-stimulated conditions (25 ng/ml, 96 h). (E-F) Quantification of FBGC formation from (D): (E) number of FBGCs per high-power field and (F) percentage of fused BMDMs. (G) Representative images of FBGCs in WT and TRPV1 KO BMDMs cultured on collagen-coated (10 μg/ml) 50 kPa PA hydrogels, with or without IL-4 plus GMCSF stimulation (25 ng/ml, 96 h). (H-I) Quantification of FBGC formation from (G): (H) number of FBGCs per high-power field and (I) percentage of fused BMDMs. Data represent n = 3 biological replicates with 5 images per group. Scale bar, 20 μm; statistical analysis by one-way ANOVA, ***p < 0.001, ****p < 0.0001.

Consistent with these findings, TRPV1-deficient BMDMs displayed marked impairments in multinucleation when cultured on rigid permanox substrates (> 1 GPa), with substantial reductions in both FBGC number (∼6-fold) and fusion index (∼15-fold) (Figure 4D-F). To directly assess the role of TRPV1 in stiffness-driven fusion, WT and TRPV1 KO BMDMs were cultured on stiff (50 kPa) hydrogels. Loss of TRPV1 significantly attenuated macrophage fusion under these conditions, resulting in ∼6-fold fewer FBGCs and an approximately 20-fold decrease in fusion index, even in the presence of IL-4 and GM-CSF (Figure 4G-I). Collectively, these findings identify TRPV1 as a key mechanosensitive regulator of macrophage multinucleation, required for efficient FBGC formation in response to both matrix stiffness and fusogenic cytokine signaling.

### TRPV1 regulates matrix stiffness-induced F-actin generation in macrophage giant cells

Cytoskeletal dynamics are central to a wide range of cellular functions, including the fusion of macrophages into multinucleated cells (49, 51-53, 61-75). Because cellular mechanical properties are largely dictated by actin polymerization, we examined whether TRPV1-dependent FBGC formation is coupled to changes in actin organization. Upon stimulation with IL-4 and GMCSF, WT BMDMs displayed prominent F-actin ring structures within FBGCs, whereas these features were absent in TRPV1-deficient cells, which also failed to form FBGCs (Figure 5A-C). To determine whether matrix rigidity influences actin remodeling through TRPV1, WT macrophages were cultured on collagen-coated PA hydrogels with defined stiffness (1 kPa vs. 50 kPa). Increased substrate stiffness markedly elevated F-actin assembly, and this response was further augmented by cytokine treatment (Figure 5D-E), suggesting a synergistic effect of mechanical and inflammatory cues. Inhibition of TRPV1 using AMG9810 suppressed this enhancement, indicating that TRPV1 is required for stiffness- and cytokine-driven actin reorganization. We next directly tested the requirement of TRPV1 under stiff conditions by comparing WT and TRPV1 KO BMDMs on 50 kPa matrices. TRPV1 deficiency significantly impaired F-actin accumulation, with approximately a fourfold reduction despite cytokine stimulation (Figure 5F-G). Together, these results establish TRPV1 as a critical mediator linking extracellular stiffness and cytokine signaling to cytoskeletal remodeling during macrophage fusion.

**Figure 5.**
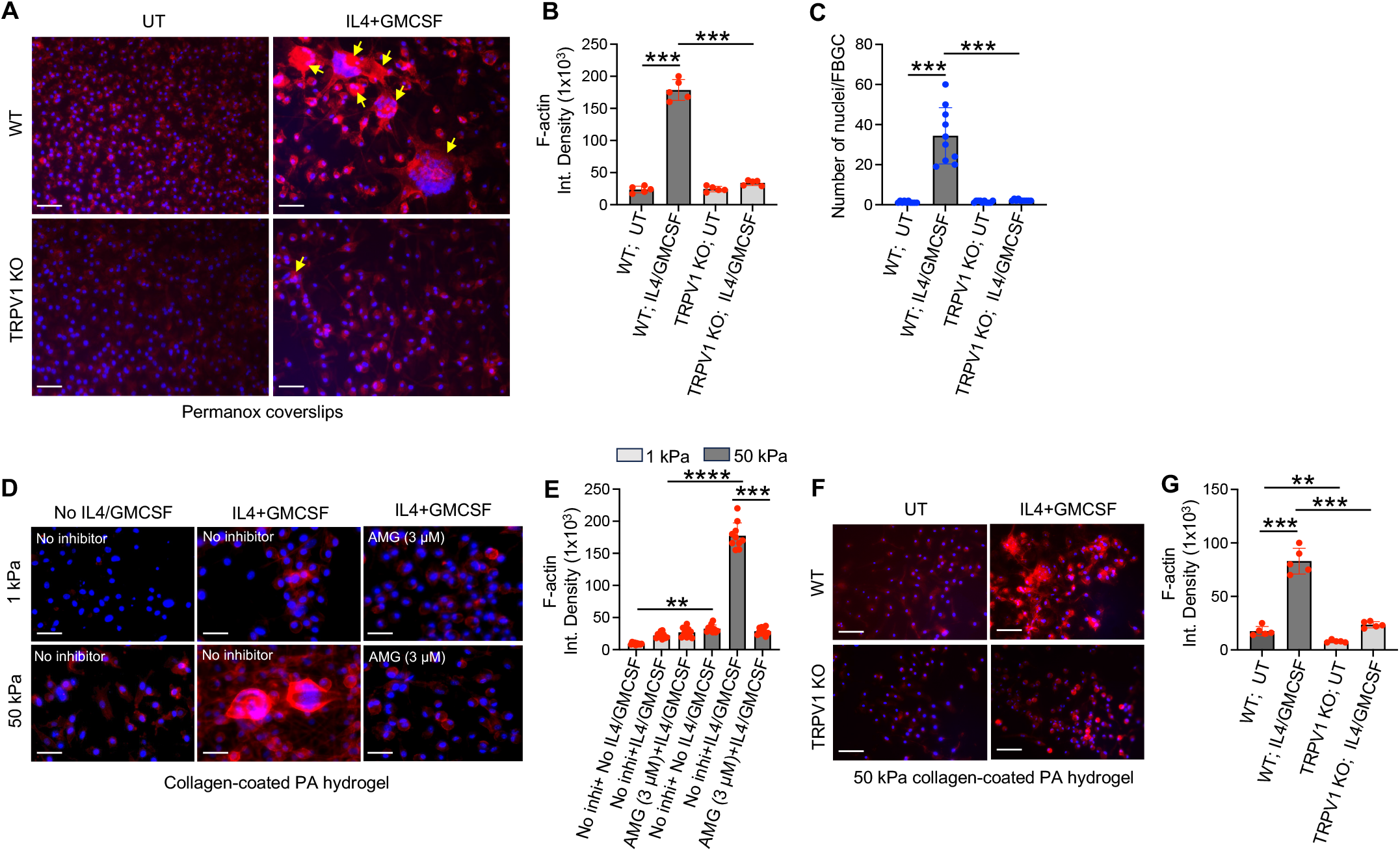
TRPV1 controls matrix stiffness-dependent F-actin assembly during macrophage giant cell formation. (A) Representative fluorescence images of WT and TRPV4 KO BMDMs, either untreated or stimulated with IL-4 plus GMCSF (25 ng/ml), showing phalloidin-labeled F-actin (red). (B) Quantification of F-actin intensity corresponding to (A). (C) FBGC formation from (A), expressed as nuclei per multinucleated cell. (D) Representative images of WT BMDMs cultured on collagen-coated (10 μg/ml) PA hydrogels of low and high stiffness, with or without IL-4 plus GMCSF stimulation, showing F-actin (red). (E) Quantification of F-actin levels from (D). (F) Representative images of WT and TRPV4 KO BMDMs cultured on 50 kPa PA hydrogels under the indicated conditions, stained for F-actin (red). (G) Quantification of F-actin levels from (F). Scale bars, 50 μm. Data in (A-G) represent 5-10 cells per group from three independent experiments. Statistical significance was determined by one-way ANOVA (**p < 0.01, ***p < 0.001, ****p < 0.0001).

### TRPV1 regulates macrophage giant cell formation independent of TRPV4 activity

Our prior work identified the mechanosensitive channel TRPV4 as a key regulator of stiffness-driven FBR and FBGC formation (48, 49, 51). To determine whether TRPV1-dependent macrophage fusion operates through TRPV4, we assessed calcium signaling in WT and TRPV1-deficient BMDMs following 48 h stimulation with IL-4 and GMCSF. Activation of TRPV1 with capsaicin elicited a robust increase in intracellular Ca^2+^ in cytokine-primed WT cells compared to untreated controls (Figure 6A-B). As anticipated, this response was markedly blunted in TRPV1 KO macrophages, confirming loss of TRPV1 function. We next examined whether TRPV4 activity is altered in the absence of TRPV1 in BMBMs. Stimulation with the TRPV4-selective agonist GSK101 induced Ca^2+^ influx in TRPV1-deficient cells (Figure 6C-D), indicating preserved TRPV4 function. Collectively, these findings demonstrate that TRPV1 loss does not impair TRPV4-mediated signaling, suggesting that TRPV1 regulates macrophage fusion independently of TRPV4 under fusogenic cytokine conditions.

**Figure 6.**
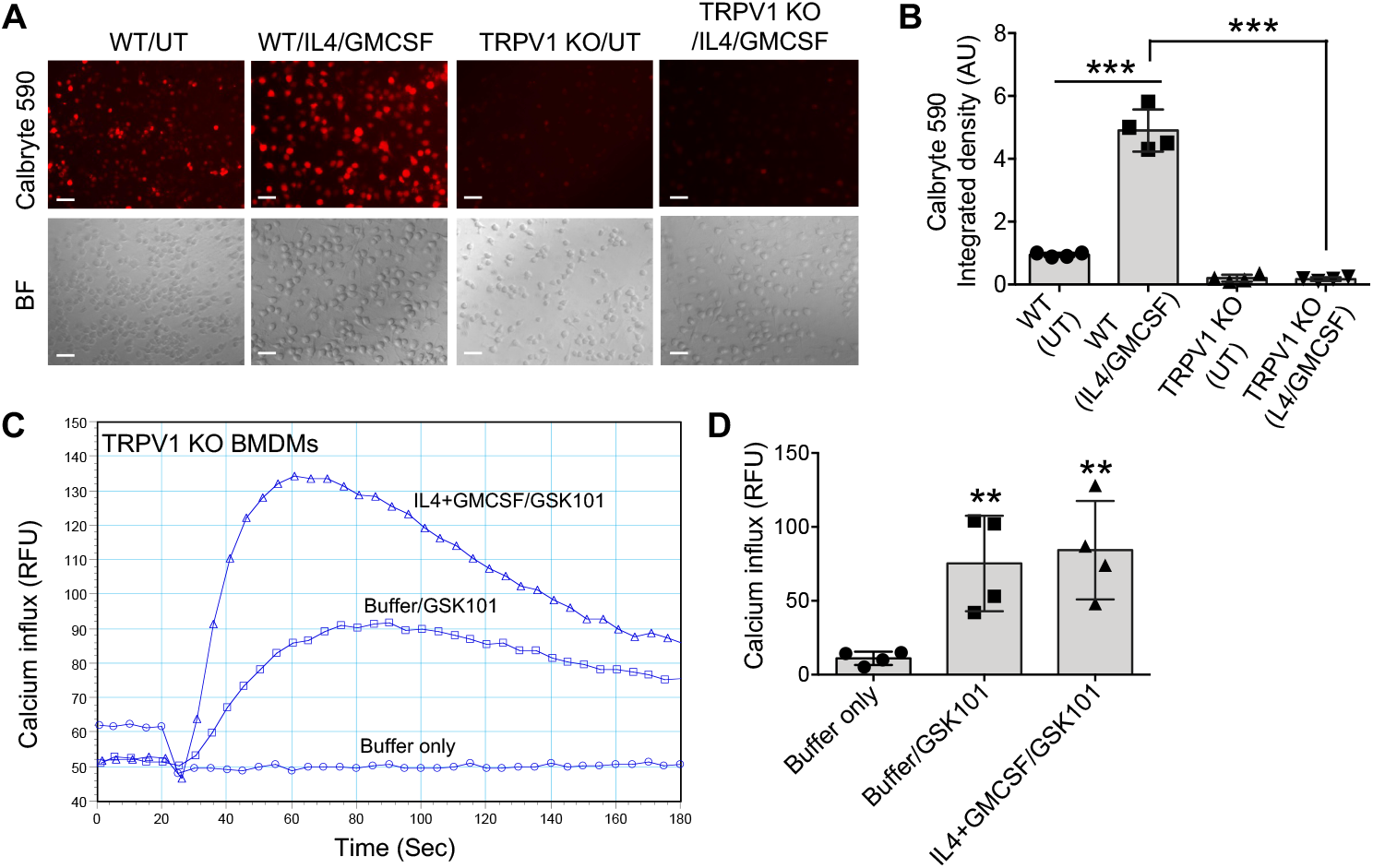
TRPV1 regulates macrophage giant cell formation independently of TRPV4 activity. (A) Spinning-disk confocal images of WT and TRPV1 KO BMDMs showing Ca^2+^ influx (red) under untreated (UT) conditions or following IL-4 plus GMCSF stimulation (25 ng/ml) in response to the TRPV1 agonist capsaicin; scale bar, 50 μm. (B) Quantification of fluorescence intensity from (A) (n = 4 fields per condition). (C) FlexStation 3 measurements of Ca^2+^ influx in TRPV1 KO BMDMs stimulated with the TRPV4-specific agonist GSK1016790A (GSK101) under untreated (buffer control) or IL-4 plus GMCSF-treated conditions (25 ng/ml). (D) Quantification of Ca^2+^ responses from (C). Experiments were performed three times with quadruplicate measurements. RFU, relative fluorescence units. Statistical significance was determined by Student’s *t*-test (**p < 0.01, ***p < 0.001).

## Discussions

In this work, TRPV1 is identified as a critical regulator of macrophage multinucleation and FBGC formation. It is expressed in bone marrow-derived macrophages and upregulated by fusogenic cytokines and inflammatory signals. Functionally, TRPV1 promotes matrix stiffness-dependent adhesion and spreading, highlighting its role in mechanosensing. It is required for efficient macrophage fusion under both cytokine-driven and stiffness-mediated conditions. Mechanistically, TRPV1 links extracellular mechanical and inflammatory cues to cytoskeletal remodeling, enabling actin reorganization necessary for fusion. Notably, TRPV1 deficiency does not affect TRPV4 activity, indicating an independent pathway. Overall, TRPV1 emerges as a novel mechanosensitive regulator of macrophage fusion, offering new insight into FBR mechanisms and a potential target to improve biomaterial biocompatibility and reduce fibrosis.

This study identifies TRPV1 as a previously unrecognized regulator of macrophage fusion and the FBR. While prior work has emphasized mechanosensitive pathways in macrophage biology - particularly TRPV4 - our findings expand this framework by demonstrating that TRPV1 integrates inflammatory and mechanical cues to control multinucleation and FBGC Formation (48, 49, 51). Rather than functioning as a redundant TRPV family member, TRPV1 emerges as a parallel, independent signaling axis that couples matrix stiffness and cytokine stimulation to cytoskeletal remodeling, a central requirement for cell fusion. Conceptually, this work advances the FBR and cell fusion fields in three keyways. First, it broadens the paradigm of macrophage mechanotransduction beyond TRPV4-centric models, introducing TRPV1 as a critical mediator of stiffness-dependent fusion. Second, it links calcium-permeable ion channel signaling directly to actin reorganization during multinucleation, providing mechanistic insight into how extracellular environments shape fusion competence. Third, it establishes a framework in which biochemical (cytokine) and biophysical (matrix stiffness) signals converge at ion channels to regulate FBGC formation, offering a more integrated view of biomaterial-immune interactions.

Despite these advances, several limitations should be noted. (i) The study relies primarily on *in vitro* system; *in vivo* validation in clinically relevant implantation models is needed to confirm the role of TRPV1 in FBR progression. (ii) While cytoskeletal remodeling is demonstrated, the downstream signaling pathways linking TRPV1-mediated Ca^2+^ influx to actin dynamics remain incompletely defined. (iii) Under specific stimuli, monocytes/macrophages fuse to form multinucleated giant cells, including Langhans giant cells (LGCs), FBGCs, and osteoclasts (). These arise via distinct cytokine cues: IFNγ drives LGCs, IL-4 plus GMCSF induces FBGCs, and RANKL plus MCSF promotes osteoclast formation. Morphologically, LGCs display polarized nuclei, FBGCs exhibit randomly distributed nuclei, and osteoclasts show scattered nuclei with foamy cytoplasm (76-78). IL-4 and GMCSF suppress osteoclastogenesis while promoting macrophage differentiation and FBGC formation (76-78). Accordingly, we used IL-4 plus GMCSF to generate FBGCs from BMDMs, identified by their characteristic random nuclear distribution.

Overall, this work positions TRPV1 as a novel therapeutic target for modulating macrophage fusion, with potential implications for improving biomaterial integration and limiting fibrosis in response to implantation.

## Acknowledgements

This work was supported by an NIH (R01EB024556) grant to Shaik O. Rahaman. We acknowledge the DLAR Imaging Core Facility, located at A.J. Clark Hall at the University of Maryland, College Park for providing imaging equipment assistance and consultation.

## Authors contributions

SOR and KRS conceived the study, designed the experiments, analyzed the data. MIK and KRS performed the experiments. SOR wrote and edited the manuscript. All authors reviewed and approved the final content of the manuscript.

## Data availability

All data generated and used during this study are included in this article.

